# Inhibition of *de novo* ceramide biosynthesis affects aging phenotype in an *in vitro* model of neuronal senescence

**DOI:** 10.1101/645879

**Authors:** Alberto Granzotto, Manuela Bomba, Vanessa Castelli, Riccardo Navarra, Noemi Massetti, Marco Onofrj, Ilaria Cicalini, Piero del Boccio, Annamaria Cimini, Daniele Piomelli, Stefano L. Sensi

## Abstract

Although aging is considered to be an unavoidable event, recent experimental evidence suggests that the process can be delayed, counteracted, if not completely interrupted. Aging is the primary risk factor for the onset and development of neurodegenerative conditions like Alzheimer’s disease, Parkinson’s disease, and Amyotrophic Lateral Sclerosis. Intracellular calcium (Ca^2+^_i_) dyshomeostasis, mitochondrial dysfunction, oxidative stress, and lipid dysregulation are critical factors that contribute to senescence-related processes. Ceramides, a class of sphingolipids involved in a wide array of biological functions, are important mediators of cellular senescence, but their role in neuronal aging is still largely unexplored.

In this study, we investigated the effects of L-cycloserine (L-CS), an inhibitor of *de novo* ceramide biosynthesis, on the aging phenotype of cortical neurons that have been maintained in culture for 22 days, a setting employed as an *in vitro* model of cellular senescence. Our findings indicate that ‘aged’ neurons display, when compared to control cultures, overt dysregulation of cytosolic and subcellular [Ca^2+^]_i_ levels, mitochondrial dysfunction, increased reactive oxygen species generation, altered synaptic activity as well as the activation of neuronal death-related molecules. Treatment with L-CS (30 µM) positively affected the senescent phenotype, a result accompanied by recovery of neuronal [Ca^2+^]_i_ signaling, and reduction of mitochondrial dysfunction and reactive oxygen species generation.

The results suggest that the *de novo* ceramide biosynthesis may represent a critical intermediate in the molecular and functional cascade leading to neuronal senescence. Our findings also identify ceramide biosynthesis inhibitors as promising pharmacological tools to decrease age-related neuronal dysfunctions.

## 1. Introduction

Aging is the time-dependent process characterized by the loss of the physiological integrity of living organisms ^1^. Although this process has been long considered to be unavoidable, recent evidence has shown that it can be delayed, if not altogether interrupted ^2^. Many factors, including environmental exposure and molecular changes, contribute to the senescence-related processes ^1,3^. The molecular effectors promote several cellular modifications like DNA damage, the activation of cell death pathways, the loss of proteostasis, cation dyshomeostasis, mitochondria dysfunction, and lipid dysregulation ^1,4–6^.

Ceramides are a class of sphingolipids involved in a wide array of biological functions including cell growth, proliferation, differentiation, and programmed cell death as well as cellular senescence ^7^. The application to cell cultures of exogenous ceramides induces, in a dose-dependent and reversible manner, the expression of senescence markers ^8^. In parallel, cellular senescence is associated with the increase of ceramide levels, a phenomenon found in age-related neurological conditions like Alzheimer’s disease (AD), Parkinson’s disease (PD), and Amyotrophic Lateral Sclerosis (ALS) patients as well as in brain aging ^9–13^.

Ceramides originate from one of two main enzyme-mediated processes: *de novo* biosynthesis from palmitoyl-CoA and serine, or cleavage of sphingolipid precursors in membranes ^14^. The former pathway involves three sequential reactions in which the enzyme serine palmitoyl transferase (SPT) serves as rate-limiting step ^14^. Importantly, pharmacological blockade of SPT activity has shown promising anti-aging and neuroprotective effects in *in vitro* and *in vivo* models of cellular senescence ^15,16^.

To gain further insights on the role played by ceramides in neuronal aging, we used L-cycloserine (L-CS), an amino acid that inhibits SPT activity both safely and selectively, and has been employed both *in vitro* and *in vivo* settings to reduce *de novo* ceramide biosynthesis ^15,17–20^. We evaluated age-driven changes occurring in young and aging neuronal cultures exposed to vehicle or L-CS, focusing on alterations in cytosolic and subcellular calcium [Ca^2+^]_i_ handling, mitochondrial functioning, spontaneous neuronal Ca^2+^ signaling, and activation of age-related and ceramide-driven molecular pathways.

## 2. Results

To examine the effects of L-CS on neuronal aging, experiments were performed in long-term culture of primary cortical neurons (hereafter termed ‘aged’ neurons) used here as an *in vitro* model of neuronal senescence ^21–23^. Cultures were maintained for 18-19 days *in vitro* (DIV) and then treated with L-CS (30 µM) or vehicle (serum-free medium) for an additional three days. At the end of treatment (at 21-22 DIV), cultures were analyzed for morphological and functional changes or harvested for further biochemical analyses (Fig. 1A). Data collected from aged neurons were compared with those obtained from vehicle- or drug-treated cultures at 11-12 DIV and assayed at 14-15 DIV (hereafter termed ‘control’ neurons). Routinely performed visual inspections of the cultures showed that aged neurons did not display overt signs of death or injury (Fig. 1B), thereby supporting the notion that the functional and biochemical changes that we have observed occurred upstream of mechanisms leading to neuronal death.

**Figure 1.**
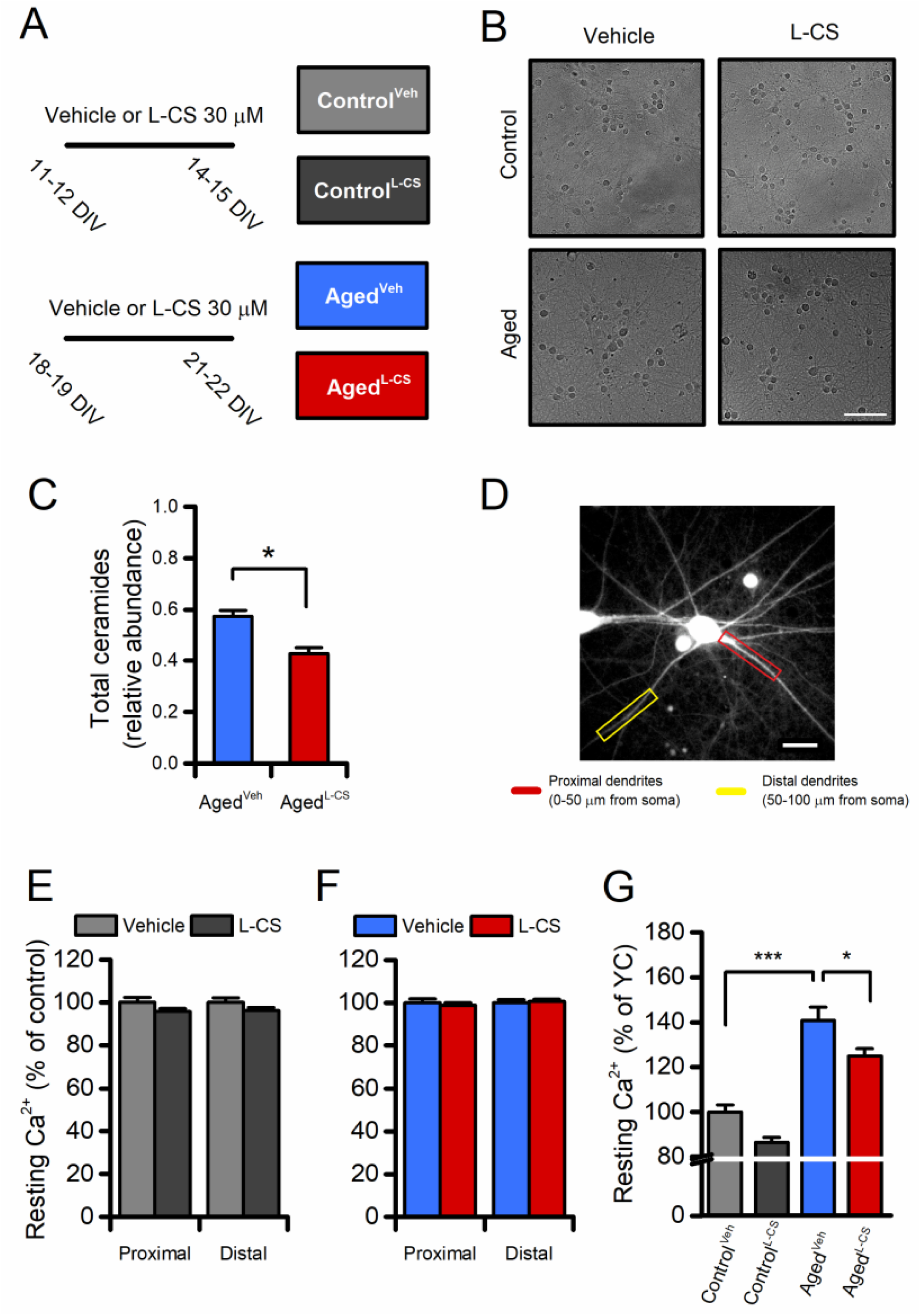
Effects of aging and L-CS on resting calcium (Ca^2+^) levels in cortical neurons. (A) The pictogram illustrates the experimental paradigm employed in the study. (B) Representative brightfield micrographs of control and aged neuronal cultures treated either with L-CS or vehicle (scale bar 100 µm). Please, note that aged cultures are devoid of signs of neuronal death. (C) Bar graphs depict the relative abundance of ceramides in vehicle- and L-CS-treated aged neurons (n=3 for both conditions). (D) Representative fluorescent micrograph of a fura-2-loaded cultured cortical neuron (the image reports dye emission when excited at 380 nm, scale bar 25 µm). (E) Bar graphs depict dendritic Ca^2+^ levels of vehicle- or L-CS-treated control neurons (vehicle: n=102 proximal and n=85 distal dendrites from 43 neurons; L-CS: n=115 proximal and n=84 distal dendrites from 38 neurons; p>0.05). (F) Bar graphs depict dendritic Ca^2+^ levels of vehicle- or L-CS-treated aged neurons (vehicle: n=182 proximal and n=156 distal dendrites from 40 neurons; L-CS: n=177 proximal and n=155 distal dendrites from 44 neurons; p>0.05). (G) Bar graphs depict somatic Ca^2+^ levels of vehicle- or L-CS-treated control and aged neurons (Control^Veh^: n=1357 cells and Control^L-CS^ n=1015; Aged^Veh^ n=539 cells and Aged^L-CS^ n=497 cells obtained from 10-23 independent experiments). In C and E-F means were compared by unpaired Student t-test. In G means were compared by two-way ANOVA followed by Tukey post-hoc test. * indicates p<0.05, *** p<0.001.

### 2.1 L-CS reduces de novo ceramide biosynthesis in aged cultured neurons

To test the drug-driven effects on ceramide biosynthesis, we quantified sphingolipids and ceramides by Liquid Chromatography/Mass Spectrometry (LC/MS-MS) in lipid extracts obtained from control and aged cultures that underwent vehicle- or L-CS-treatment (Fig. 1A, B). Analysis of the lipid profiles indicates that aged neurons respond to the 3-day L-CS treatment with a significant reduction in the levels of total ceramides (d18:1/16:0, d18:1/18:0, d18:1/22:0, and d18:1/24:0; Fig.1C) and sphinganine (Supplementary Fig. 1). Aged cultures also showed a trend toward a statistically significant increase in ceramide levels when compared to control neurons (Supplementary Fig. 1). Of note, the ceramide reduction appears to be mainly driven by inhibition of the *de novo* pathway as sphinganine (an intermediate of the *de novo* pathway), but not sphingosine (which can generate ceramide through sphingomyelin hydrolysis), was found to be affected by the L-CS treatment (Supplementary Fig. 1).

### 2.1 L-CS reduces resting Ca^2+^ levels in aged neurons

The effects of 3-day L-CS or vehicle treatment on [Ca^2+^]_i_ were measured by using the high-affinity (K_d_≈140nM) ratiometric dye fura-2. The analysis of resting fura-2 signals showed that no regional differences in [Ca^2+^]_i_ levels were detectable in proximal or distal dendrites of aged or control neurons treated with either vehicle or L-CS (Fig. 1D-F). By contrast, L-CS treatment caused a reduction in resting somatic [Ca^2+^]_i_ levels in aged neurons (Fig. 1G). Of note, a Z≈40% increase in resting [Ca^2+^]_i_ was observed when comparing vehicle-treated control and aged neurons, thereby lending support to the notion that our *in vitro* senescence model exhibits signs of age-dependent Ca^2+^ dyshomeostasis ^5,24^. No significant differences were observed when comparing vehicle-and drug-treated control neurons (Fig. 1G).

### 2.2 L-CS treatment marginally affects main neuronal Ca^2+^ stores

Mitochondria and the endoplasmic reticulum (ER) constitute the main intracellular Ca^2+^ stores. To test whether Ca^2+^ accumulation in these organelles is affected by L-CS treatment, fura-2-loaded control and aged neurons were exposed to 5 µM carbonyl cyanide 3-chlorophenylhydrazone (CCCP, a mitochondrial uncoupler that collapses the mitochondrial membrane potential, Δp) or 10 µM cyclopiazonic acid [CPA, a sarco/endoplasmic reticulum Ca^2+^-ATPase (SERCA) inhibitor], two agent employed to promote cation release from mitochondria or the ER, respectively. Analysis of the cytosolic [Ca^2+^]_i_ changes showed that exposure to L-CS did not affect mitochondrial Ca^2+^ release (Fig. 2A-C). A significant increase in mitochondrial Ca^2+^ release was instead found in aged neurons (Fig. 2A-C). As our cultures showed modest and inconsistent CPA-dependent ER-Ca^2+^release (data not shown) and to facilitate the evaluation of this pathway, ER was pre-loaded with Ca^2+^ derived from the opening of the voltage-gated Ca^2+^ channels (VGCCs) as result of a 5-min depolarization induced by exposing cultures to 60 mM K^+^. Compared to vehicle-treated and age-matched neurons, Ca^2+^ released from the ER was found to be lower in L-CS-treated control cultures (Fig. 2D-F). No differences were observed when comparing vehicle-treated control neurons with vehicle- and L-CS-treated aged cultures (Fig. 2D-F).

**Figure 2.**
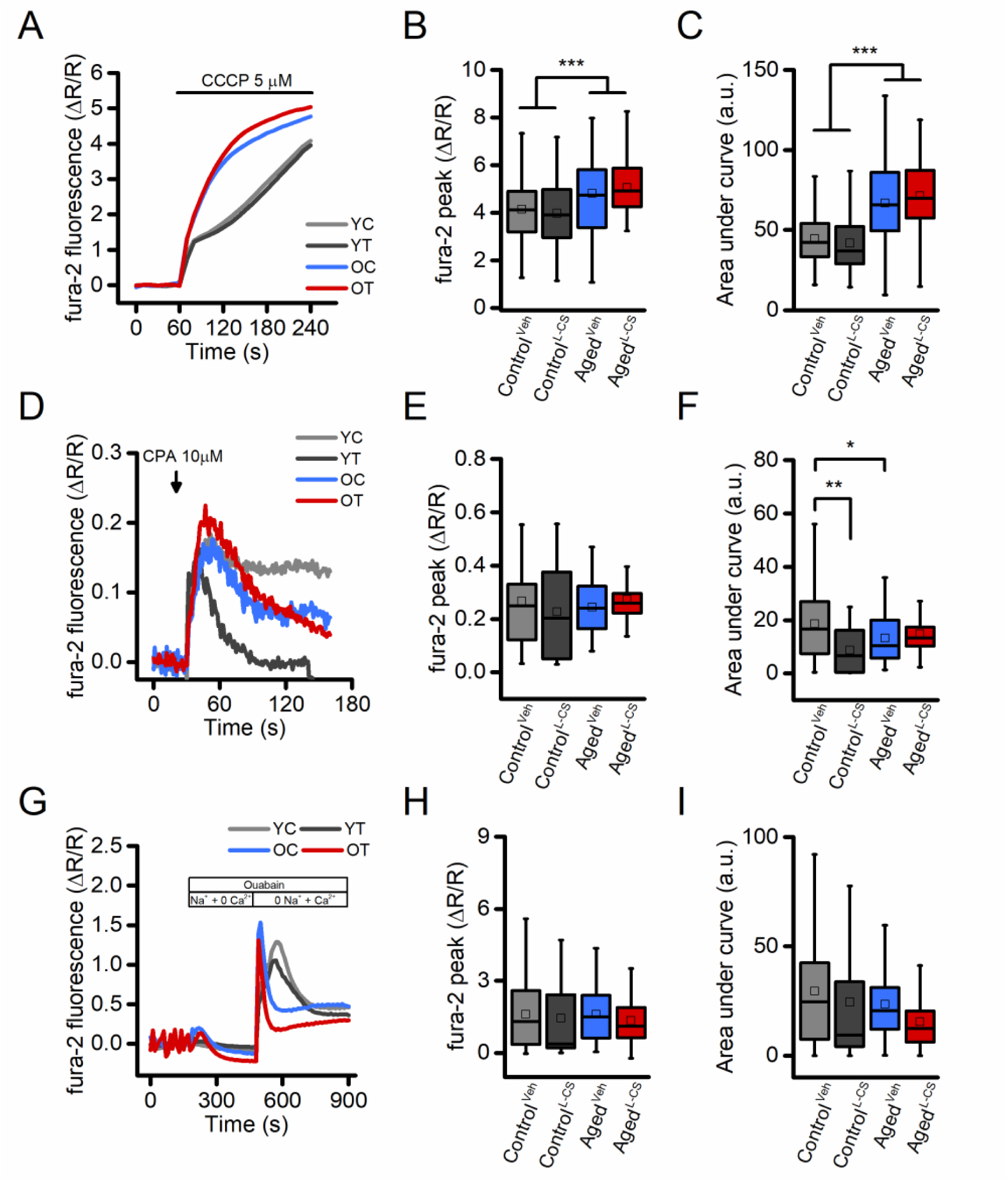
Effects of aging and L-CS on intracellular Ca^2+^ stores and NCX activity in cortical neurons. (A) Time course of CCCP-stimulated Ca^2+^ release from mitochondria. Traces represent the average response to a 3 min exposure to 5 µM CCCP (Control^Veh^: n=226 cells and Control^L-CS^ n=140; Aged^Veh^ n=91 cells and Aged^L-CS^ n=67 cells obtained from 7-19 independent experiments). (B) Box plots depict Ca^2+^ peak obtained in the four study groups. (C) Box plots depict Ca^2+^ changes expressed as AUC (a.u.). (D) Time course of CPA-stimulated Ca^2+^ release from the ER. Traces represent the average response to a 2 min exposure to 10 µM CPA (Control^Veh^: n=50 cells and Control^L-CS^ n=33; Aged^Veh^ n=54 cells and Aged^L-CS^ n=48 cells obtained from 3-4 independent experiments). (E) Box plots depict Ca^2+^ peak obtained in the four study groups. (F) Box plots depict Ca^2+^ changes expressed as AUC (a.u.). (G) Time course of NCX activity imaged by stimulating exchanger reverse operational mode (Control^Veh^: n=163 cells and Control^L-CS^ n=122; Aged^Veh^ n=106 cells and Aged^L-CS^ n=98 cells obtained from 2 independent experiments). (B) Box plots depict Ca^2+^ peak obtained in the four study groups. (C) Box plots depict Ca^2+^ changes expressed as AUC (a.u.). Means were compared by two-way ANOVA followed by Tukey post-hoc test. * indicates p<0.05, *** p<0.001.

### 2.3 L-CS treatment does not modify NCX activity

Ceramides can modulate the activity of the plasmalemmal Na^+^-Ca^2+^ exchanger (NCX) ^25^, a low capacitance high-affinity system that critically regulates cellular [Ca^2+^]_i_. We, therefore, investigated the effects of ceramide on NCX functioning in L-CS or vehicle-treated cultures. Data obtained from this set of experiments were compared with the ones originated from age-matched and vehicle-treated sister cultures. In the experiments, fura-2 loaded neurons were exposed to a Ca^2+^ free medium while in the presence of the Na^+^/K^+^-ATPase pump blocker, ouabain (100 µM), a maneuver set to promote the intracellular accumulation of Na^+^. Ouabain-treated cultures were then switched to a Na^+^-free medium to force the NCX to operate in the reverse mode to promote Ca^2+^ influx. Analysis of the cytosolic [Ca^2+^]_i_ rises showed that L-CS does not affect NCX activity in control and aged cultures treated with either vehicle- or L-CS (Fig. 2G-H). Changes in NCX functioning were observed; however, when comparing [Ca^2+^]_i_ rises in control and aged cultures (Fig. 2G-H).

### 2.4 L-CS treatment reduces age-driven mitochondrial dysfunction

Mitochondrial functioning is a key target of the aging process ^2,21^. In our experimental setting, effects on the organelle membrane potential were assessed by employing tetramethyl rhodamine ethyl ester (TMRE), a mitochondrial probe sensitive to changes of Δp. TMRE-loaded neurons were imaged for up to 60 seconds under resting conditions and after exposure to 10 µM CCCP, a maneuver that promotes a rapid and complete loss of Δp. Data from this set of experiments indicate that aging neurons display signs of mitochondria dysfunction when compared to control cultures. L-CS treatment was effective in reducing the Δp loss in aging neurons (Fig. 3A-C). A modest (≈10%) increase was also observed in L-CS-treated control neurons when compared with age-matched cultures (Fig. 3A-C). In a subset of experiments, we also evaluated age-dependent changes in mitochondrial morphology of the cultured neurons. To that aim, neurons loaded with Mitotracker, a Δp-independent mitochondrial stain, were imaged with super-resolution confocal microscopy and analyzed with the Mitochondrial Network Analysis (MiNA) toolkit ^26^. Surprisingly, no significant age-dependent morphological changes were found (Fig. 3D and Supplementary Table 1).

**Figure 3.**
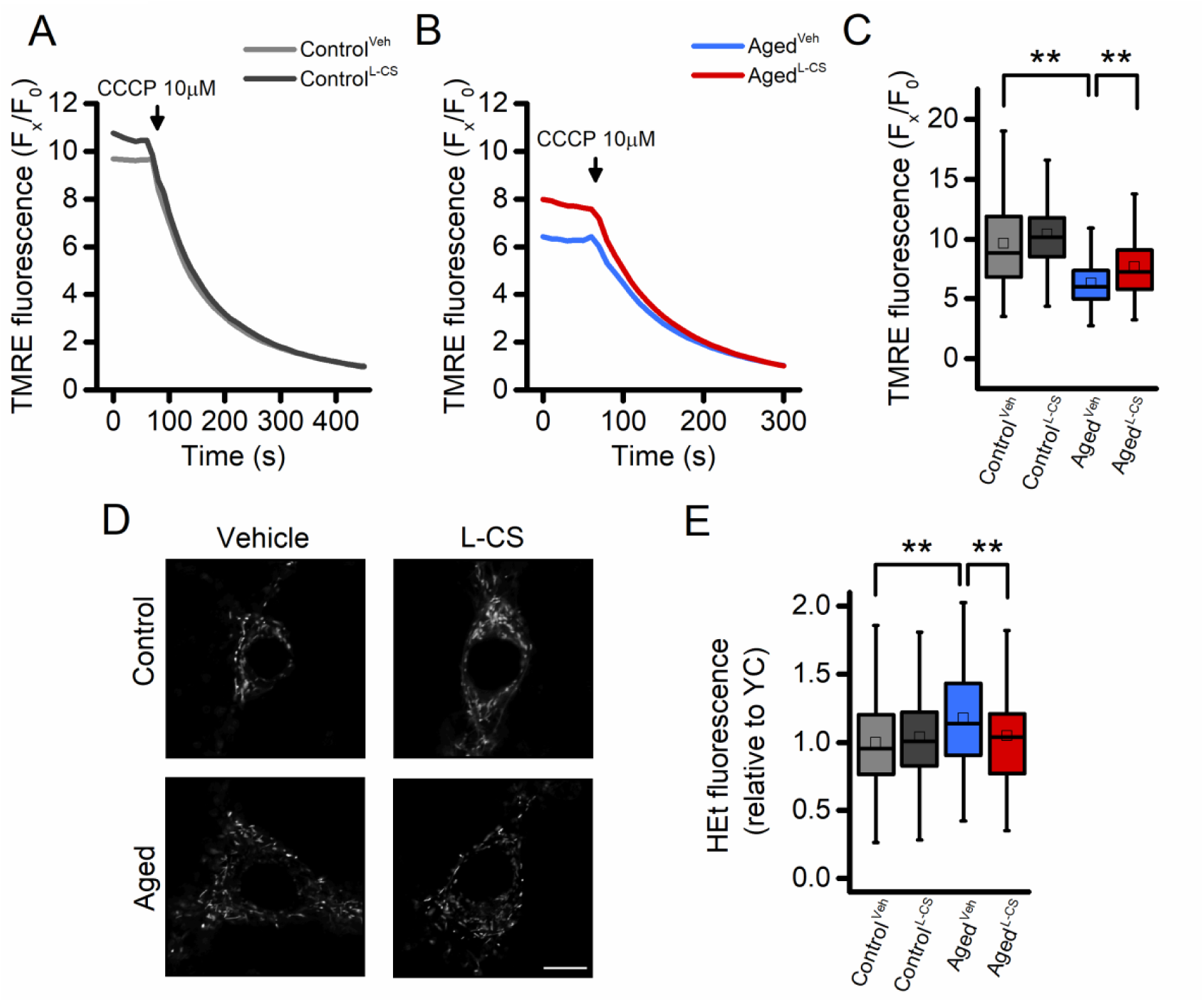
Effects of aging and L-CS on mitochondrial functioning, morphology, and ROS generation in cortical neurons. (A-B) Time course of CCCP-driven dissipation of the mitochondrial Δp. Traces represent the average response to 10 µM CCCP exposure. (Control^Veh^: n=189 cells and Control^L-CS^ n=228; Aged^Veh^ n=255 cells and Aged^L-CS^ n=240 cells obtained from 4-5 independent experiments). Please, note that aged cortical cultures require a shorter CCCP exposure time (4 min) to reach resting fluorescence levels (B). (C) Box plots depict quantification of data shown in A and B. (D) Representative super-resolution confocal images of Mitotracker Green-loaded control and aged neuronal cultures treated either with L-CS or vehicle (for quantification see supplementary table 1, n=4-6 neurons per condition; scale bar 10 µm). Please, note that no major morphological changes were observed among study groups. (E) Box plots depict normalized resting HEt fluorescence obtained from the four study groups (Control^Veh^: n=361 cells and Control^L-CS^ n=332; Aged^Veh^ n=233 cells and Aged^L-CS^ n=301 cells obtained from 5-9 independent experiments). Means were compared by two-way ANOVA followed by Tukey post-hoc test. * * indicates p<0.01.

### 2.5 L-CS treatment reduces reactive oxygen species (ROS) generation in aged cultures

Aberrant ROS generation and accumulation of ROS-driven by-products are key features of aging ^27^. To assess the effect of L-CS treatment on ROS production, we employed the probe hydroethidine (HEt). Cortical cultures were loaded with HEt and basal fluorescence signals recorded. The results show that aged neurons exhibit increased ROS production, compared to controls (Fig. 3E). L-CS administration abrogated the effects of age (Fig. 3E).

### 2.6 L-CS treatment reduces age-driven neuronal hyperexcitability in vitro

To assess whether L-CS treatment affects spontaneous [Ca^2+^]_i_ transients, an index of neurodegenerative hyperexcitability, we employed high-speed real-time microfluorimetric Ca^2+^ imaging ^28^. Cultures were loaded with the single wavelength, high-affinity (K_d_ ≈ 335 nM) and Ca^2+^ sensitive fluorescent probe fluo-4 and spontaneous changes in somatic [Ca^2+^]_i_ were evaluated in terms of spike frequency and amplitude. In line with previous studies ^29^, aged neurons showed an increased number of transients per minute, coupled with decreased Ca^2+^ spike amplitude, when compared with control cells (Fig. 4A-D). Compared with age-matched sister cultures, L-CS treatment resulted in a significant reduction of spike frequency and increased transient amplitudes in both control and aged cultures (Fig. 4A-D). These results in combination with the altered levels of [Ca^2+^]_i_ found in the soma of aged neurons left open the possibility that the changes in somatic [Ca^2+^]_i_ transients might be due to defective intracellular cation handling or to increased synaptic activity. To test these possibilities, we evaluated Ca^2+^ transients in dendrites as these compartments did not exhibit signs of Ca^2+^ dysregulation (Fig. 1E-F). Dendrites of vehicle- and L-CS-treated aged cultures did not show significant changes in spontaneous Ca^2+^ transients (Fig. 4 E-H), thereby suggesting that the altered somatic [Ca^2+^]_i_ rises are primarily due to defective Ca^2+^ homeostasis. Of note, L-CS treatment was more effective in reducing spike frequency in control neurons compared to aged cells (Fig. 4C). Similar differences in somatic Ca^2+^_i_ transients were obtained when replicating the experiments with the slower (but ratiometric) dye fura-2 (data not shown). Overall, these findings indicate that L-CS treatment reduces the altered somatic Ca^2+^ signaling driven by spontaneous synaptic activity.

**Figure 4.**
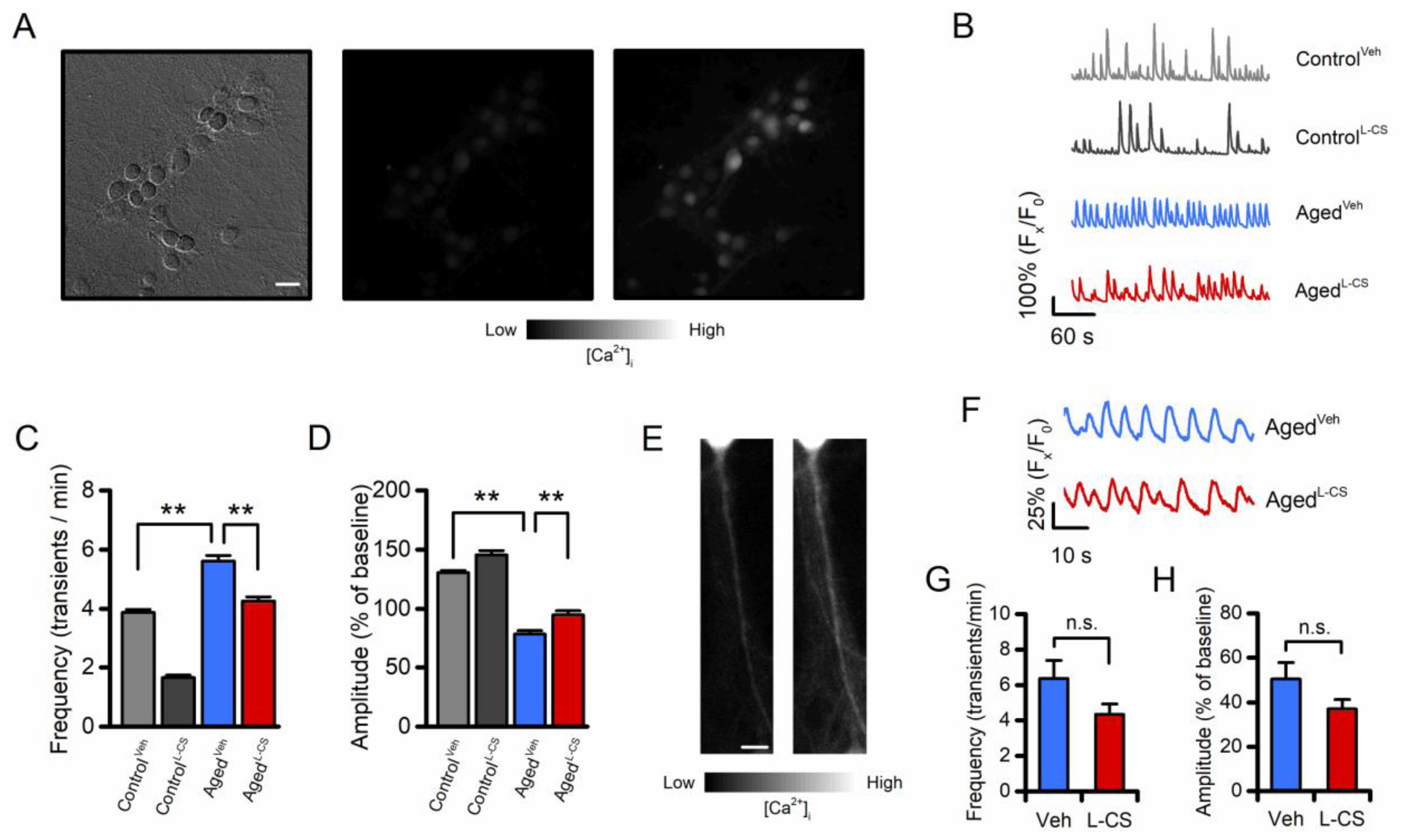
Effects of aging and L-CS on Ca^2+^ transient frequency and amplitude. (A) Representative brightfield (left) and fluorescent (middle and right) micrographs of a fluo-4-loaded aged^L-CS^ neuronal culture employed to monitor spontaneous Ca^2+^ transients (scale bar 25 µm). Greyscale fluorescent images show cortical neurons before (middle) and during (right) a Ca^2+^ transient. (B) Time course of somatic spontaneous Ca^2+^ oscillations in the four study groups. Each trace depicts a single neuron representative of at least three independent experiments. (C) Bar graphs depict average transient frequencies of vehicle- or L-CS-treated control and aged neurons (Control^Veh^: n=499, Control^L-CS^ n=253, Aged^Veh^ n=367, and Aged^L-CS^ n=293 cells obtained from 15-38 experiments). (D) Bar graphs depict the average Ca^2+^ transient amplitude in the four study groups [samples are the same as in (C)]. (E) Representative greyscale fluorescent micrographs of a fluo-4-loaded primary dendrite before (left) and during (right) a Ca^2+^ transient. (F) Time course of dendritic spontaneous Ca^2+^ oscillations in the Aged^Veh^ and Aged^L-CS^ cultured neurons. Each trace depicts a single dendrite representative of at least three independent experiments. (G) Bar graphs depict average transient frequencies of Aged^Veh^ and Aged^L-CS^ dendrites (Aged^Veh^ n=21 and Aged^L-CS^ n=27 dendrites from 12-18 experiments). (H) Bar graphs depict the average dendritic Ca^2+^ transient amplitude in the four study groups [samples are the same as in (G)]. In C and D means were compared by two-way ANOVA followed by Tukey post-hoc test. In G and H means were compared by unpaired Student t-test. ** indicates p<0.01, n.s. indicates not significant.

### 2.7 L-CS treatment reduces the age-related markers of neuronal demise

To test whether L-CS affects the molecular markers that modulate neuronal functioning, western blot (WB) analyses were performed on cell lysates from vehicle- and drug-treated cultures. When compared to control cultures, WB analysis of aged neurons showed increased expression levels of p53 and increased phosphorylation, and thus activation, of JNK, two molecules implicated in the regulation of cellular senescence programs (Fig. 5A-B) ^30,31^. These age-driven changes were reverted by L-CS treatment (Fig. 5A-B). The drug also promoted a reduction of pP38 levels, a kinase associated with senescence (Fig. 5C). When analyzed by ANOVA, the changes showed a trend towards significance (Fig. 5C; Tukey post-hoc test, P=0.06) and reached statistical significance when not corrected for multiple comparisons (Fisher LSD post-hoc test, P=0.01). No drug-related effects were observed in the control cultures (Fig. 5C). In parallel, we investigated age- and drug-related effects on the active forms of AKT and ERK5 (pAKT and pERK5, respectively), two kinases involved in neuronal survival ^32,33^. L-CS treatment was effective in reverting age-driven down-regulation of pERK5 and pAKT (Fig. 5D-E). The drug also promoted increased pAKT levels in control cultures (Fig. 5D). Overall, these findings indicate that L-CS treatment decreases the levels of senescence-associated markers and, at the same time, corrects age-driven loss of pro-survival molecules.

**Figure 5.**
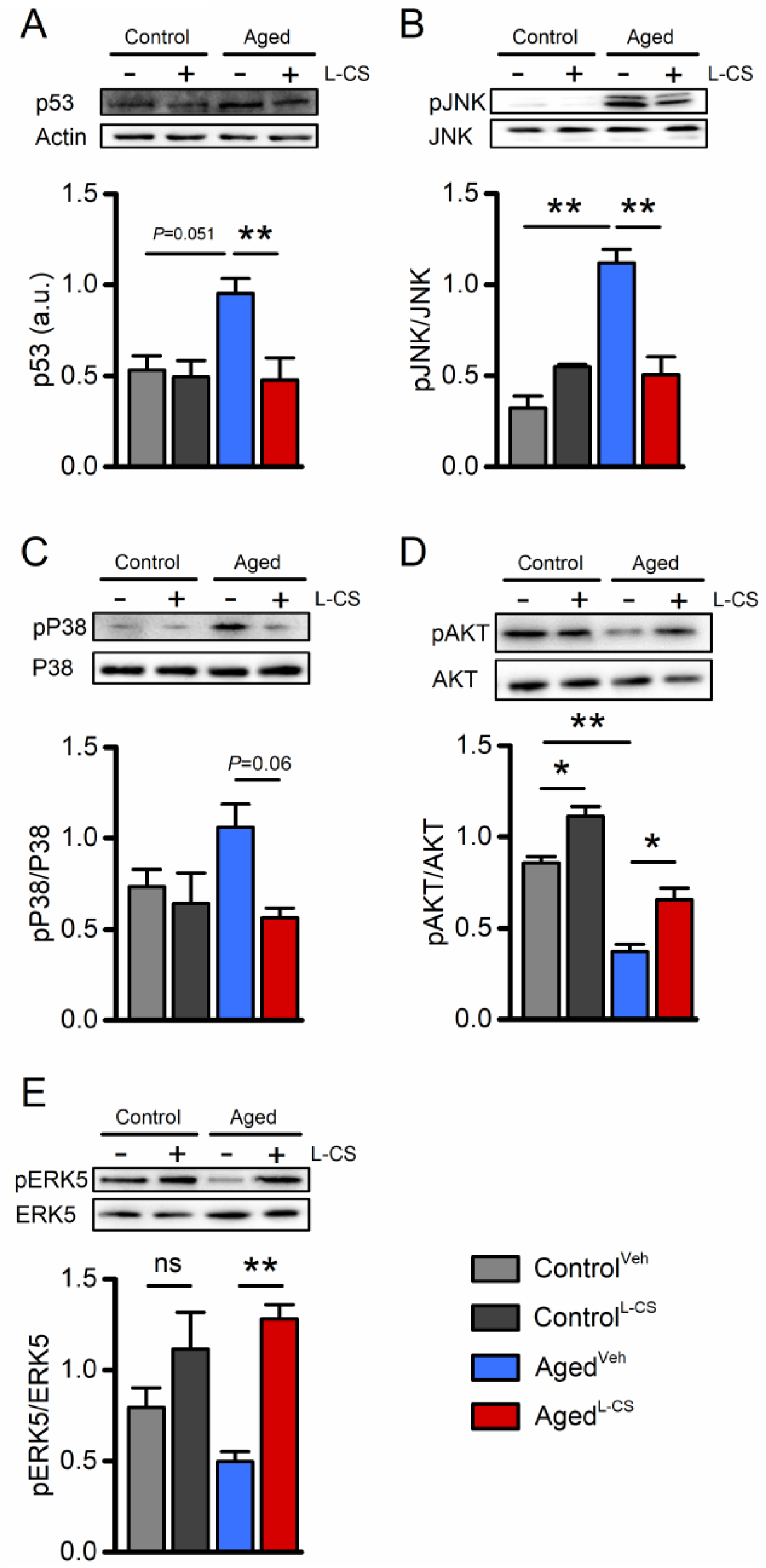
Effects of aging and L-CS on senescence-related molecular markers. Western blots show L-CS-or vehicle-driven effects on senescence-associated markers obtained from protein extracts of control and aged cortical cultures. Each image is representative of three independent experiments. (A) Bar graphs depict p53 levels in the four study groups (n=3). (B) Bar graphs depict pJNK levels in the four study groups (n=3). (C) Bar graphs depict pP38 levels in the four study groups (n=3). (D) Bar graphs depict pAKT levels in the four study groups (n=3). (E) Bar graphs depict pERK5 levels in the four study groups (n=3). Means were compared by two-way ANOVA followed by Tukey post-hoc test. * indicates p<0.05, ** indicates p003C0.01, n.s. indicates not significant.

## 3. Discussion

In this study, we used an *in vitro* model to characterize the effects of pharmacological blockade of *de novo* ceramide biosynthesis on neuronal senescence. The results indicate that exposure to the SPT inhibitor, L-CS, concurrently lowers ceramide levels in aged neurons and attenuates critical molecular and functional changes induced by aging. In particular, we found that L-CS actively counteracts the age-related mitochondrial dysfunction and neuronal Ca^2+^ dyshomeostasis.

### Age-related functional and molecular changes on cultured cortical neurons

Long-term culturing of cortical neurons promoted significant changes in cytosolic and subcellular [Ca^2+^]_i_ levels (Figs. 1 and 2). These findings are in line with and lend further support to the “Ca^2+^ hypothesis of brain aging” ^24,34,35^. This model posits that changes in the mechanisms that regulate [Ca^2+^]_i_ homeostasis play a pivotal role in the expression of physiological and pathological aging ^35^. Our findings extend this notion and indicate that age-driven [Ca^2+^]_i_ dysregulation extend to and differentially affects subcellular compartments.

A large body of evidence indicates that mitochondrial dysfunction is a crucial regulator of cellular senescence and age-related pathologies ^1,36,37^. In agreement with this notion, our aged neurons show overt signs of mitochondrial Ca^2+^ accumulation and organelle dysfunction, as indicated by the presence of significant Δp loss when compared to control cultures (Fig. 3). Unexpectedly, no significant changes in mitochondrial morphology, a proxy of the organelle and cellular health, were found in the aged neurons (Fig. 3D). A possible explanation for this lack of effect might be found in work indicating that functional alterations often precede the appearance of morphological changes ^38^. Mitochondria are also the primary generator of ROS (Supplementary Fig. 2) and compelling evidence indicates that the organelle dysfunction participates in the build-up of oxidative stress that occurs upon aging ^1^. In line with this notion, we found that aged neurons show a modest (≈20%) but significant increase in ROS production when compared to control neurons at rest (Fig. 3E).

No changes were observed in the amount of ER-Ca^2+^ release when comparing aging and control neurons (Fig. 2D-E), thereby ruling out a significant contribution exerted by ER stress on the senescent phenotype of our cells. Similarly, when comparing aged and control cultures, no significant changes were observed in the overall activity of the NCX (Fig. 2F-H). However, age-driven alterations in the NCX timing of activation were observed (Fig. 2F), a modification that may contribute to the [Ca^2+^]_i_ alterations observed in our model ^39^.

Growing evidence indicates that neuronal hyperexcitability is a crucial early feature of aging-related conditions, such as AD ^28,40,41^. Previous findings from our and other laboratories have shown that neuronal cultures display patterns of spontaneous Ca^2+^ oscillations that mirror the neuronal firing status and the *in vivo* phenotype ^28,40,41^. The analysis of this spontaneous synaptic activity indicates that aged neurons exhibit an increased frequency of [Ca^2+^]_i_ transients along with decreased spike amplitudes, two indirect signs of hyperexcitability ^28,41^. These findings parallel *in vivo* observations in hippocampal and cortical neurons of clinical and preclinical models of physio-pathological aging ^29,42–44^.

Along with functional changes, aged cortical neurons showed alterations in molecular hallmarks associated with neuronal senescence ^45–47^. We found that aged neurons exhibit higher expression levels of p53 and increased activation of pJNK and pP38, three molecules implicated in neuronal demise ^48,49^. In parallel, we observed an age-dependent decrease in the levels of the pro-survival kinases pAKT and pERK5 ^32,50^.

### Functional and molecular effects of L-CS on aged neurons

L-CS supplementation is largely employed in *in vitro* and *in vivo* settings to promote a robust decrease in ceramide levels through the inhibition of the *de novo* biosynthetic pathway ^15,19,20,51–54^. In line with this notion, L-CS was effective in reducing the ceramide pool in our aged neurons when compared to vehicle-treated cells. A modest drug-related effect was also observed in control cells. This discrepancy can be explained by previous findings indicating that the *de novo* pathway plays a primary role in ceramide biosynthesis during senescence ^15^, a mechanism likely occluded in our control cultures. Ceramide accumulation, following acute stressful challenges, participates in the activation of apoptosis in several cellular systems, including neurons. Given the central role of mitochondria in the activation of apoptotic signals, it is therefore conceivable that the L-CS-driven reduction in ceramide levels may also affect the organelle functioning. In line with this notion, we found a robust recovery (≈50%) of the mitochondrial Δp and a reduction of the ROS production in L-CS-treated neurons when compared to vehicle-treated aged-matched cultures. These findings are also in line with previous reports showing that ceramides contribute to mitochondrial dysfunction by impairing the organelle electron transport chain, generating ROS, and promoting the permeabilization of the mitochondrial outer membrane ^55,56^. ROS also contribute to the activation of ceramide-releasing enzymes ^57^, thereby suggesting the presence of a feed-forward loop in which ceramide accumulation, mitochondria impairment, and oxidative stress act synergistically to promote neuronal impairment.

To investigate the downstream effectors of the mitochondrial pathways involved in neuronal dysfunction, we evaluated changes in levels of p53, pJNK, and pP38. The selection of these proteins was driven by the fact that 1) these factors are altered in and participate to senescence-related processes; 2) their activity is intertwined with mitochondrial dysfunction; 3) their levels are modulated by ceramides ^15,48,58–60^. L-CS was found to be effective in reducing the increased levels of p53, pJNK, and pP38 (Fig. 5) shown by the aging cultures. These results support the idea that L-CS, by reducing *de novo* ceramide biosynthesis, prevents the mitochondria-driven activation of senescent-related pathways.

Our findings converge towards the possibility of a beneficial effect of L-CS in aging cultures. The analysis of the data on spontaneous synaptic activity supports this hypothesis and indicates that L-CS is effective in reducing the aberrant neuronal firing of aged cultures (Fig. 4). Thus, one can envision that the age-dependent accumulation of ceramide adducts is a critical contributor to the functional alterations in neuronal connectivity observed in either preclinical models or neurological conditions ^43,61,62^. A robust drug-related effect was also observed in control cultures, a setting that exhibited only modest reductions of ceramide levels (Fig. 5). This discrepancy can be, at least in part, explained by drug-related effects on the *in vitro* neuronal development ^7^ and/or by off-target effects of the compound ^63^.

### Conclusions

The results presented here provide experimental support for the presence of an aging-related cascade of events that include mitochondrial dysfunction, Ca^2+^ dysregulation, impaired neuronal Ca^2+^ signaling, and alterations of aging-related markers in which *de novo* ceramide biosynthesis acts a critical intermediate. Our findings indicate that ceramide modulation is not a mere response to aging-related stimuli but may play instead an active role in shaping the senescence-related processes ^13^. Indeed, they are in line with a growing body of literature supporting a key role for lipids and lipid dysmetabolism in aging and neurodegenerative conditions ^12,13,64–69^. In this context, ceramides are emerging as important peripheral biomarkers of age-related pathologies since changes in their plasmatic levels are now considered to be highly predictive of aging-related cognitive decline and conversion to AD ^13,70,71^.

The present study has several limitations. For instance, our near-pure neuronal system does not investigate the impact of ceramides on the senescent phenotype of non-neuronal cells (i.e., astrocytes, microglia, etc.), an important area of investigation given the key role played by ceramides in promoting the glia-mediated release of pro-inflammatory cytokines in the context of cell death ^16,52,72^. Furthermore, due to the limitations of our experimental setting, we have not explored sex-related differences as gender has been recently shown to have a significant impact on ceramide metabolism ^66^. Finally, the L-CS-driven effects occasionally observed in control cultures suggest that the compound may also act through mechanisms independent on ceramide biosynthesis, which warrant further investigation. Although the L-CS biological activity is distinct from the one modulated by D-cycloserine (an N-methyl-D-aspartate receptor co-agonist), it cannot be ruled out that a marginal isomerization of the compound accounts for some of the observed off-target effects.

Our results raise several intriguing questions on the interplay between ceramides and other neurodegenerative markers. For instance, do ceramides act on the same targets affected by proteins and pathways involved in AD, PD, or ALS (i.e., amyloid, tau, α-synuclein, TDP-43) or do they engage independent and/or synergistic pathways? Is there a common molecular trigger for the accumulation of ceramides and misfolded proteins? Does the manipulation of the pathological accumulation of amyloid, tau, α-synuclein, or TDP-43 affect ceramide metabolism or vice versa? Testing these questions in disease-relevant preclinical and clinical settings will have a great translational value. Nevertheless, the present results indicate that the pharmacological modulation of the *de novo* ceramide biosynthesis may be a promising target for the treatment of age-related pathologies as shown in preclinical models of AD ^20^, insulin resistance ^51^, and microglia-driven inflammation ^52^, conditions in which the ceramide build-up has been proposed to be involved in the disease onset and progression. Also, the L-CS off-target effects and its short half-life offer the possibility to investigate and test novel, brain penetrant, long-lasting, and more selective SPT inhibitors ^16^.

## 4. Experimental procedures

### 4.1 Materials

Culture media and sera were purchased from GIBCO (ThermoFisher). All the fluorescent indicators employed in the study (fura-2 AM, fluo-4 AM, hydroethidine, TMRE, and MitotrackerGreen FM) were obtained from Molecular Probes (ThermoFisher). Unless otherwise specified, all commonly used chemicals were from Sigma-Aldrich.

### 4.2 Neuronal cortical cultures

The procedures involving animals were approved by the institutional Ethics Committee (47/2011/CEISA/COM) and performed following institutional guidelines and national and international laws and policies. Female mice were caged in groups, while male mice were single-housed. Mice were kept on a 12:12 light/dark cycle and had free access to food and water. All efforts were made to minimize the number of animals employed and their suffering. Neuronal cortical cultures were, as previously described ^41^, prepared from fetal CD1 mice at 14 days of gestation and plated onto Poly-DL-lysine (100 µg/ml) and laminin (5 µg/ml) coated Petri dishes or glass coverslips. Three days after plating, the proliferation of non-neuronal cells was halted by the addition of cytosine β-D-arabinofuranoside (5 µM). Every three days 25% of the growth medium was replaced with fresh Neurobasal. Routinely-performed functional experiments and visual inspections show that the cytostatic treatment robustly affected glia proliferation (around 5–10 % of non-neuronal cells per culture).

### 4.2 Ceramide quantification

Ceramides analysis was performed as previously described ^73^. Briefly, 100 µL of sample, lysed with a probe sonicator, were mixed to 300 µL of chloroform:methanol 2:1 v/v with an internal standard mix containing sphinganine d17:0, sphingosine d17:1, sphingosine-1-phosphate d17:1, sphinganine-1-phosphate d17:0, sphingosine-1-phosphate d17:1, glucosylceramide (d18:1/17:0), and ceramide (d18:1/17:0). The organic phase was dried and reconstituted in 100 µL of H_2_O:methanol:isopropanol:acetonitrile (7:2:0.5:0.5, v/v). The LC-MS/MS system was an HPLC Alliance HT 2795 Separations Module coupled to Quattro UltimaPt ESI tandem quadrupole mass spectrometer (Waters Corporation) operating in the positive ion mode. For the chromatographic separation, Ascentis Express Fused-Core C18 2.7 µm, 7.5 cm × 2.1 mm columns were used. Elution was achieved through a gradient of mobile phases, starting from 50% to 100% of methanol:isopropanol:acetonitrile 4:1:1 v/v (solvent B), water was used as solvent A. The total run time was 25 min. The flow rate was 0.25 mL/min. The capillary voltage was 3.8 kV, source temperature was 120 °C, the de-solvation temperature was 400 °C, and the collision cell gas pressure was 3.62×10-3 mbar argon. Chromatograms were used quantify the following molecules: sphinganine (d18:0), sphingosine (d18:1), sphinganine-1-phosphate (d18:0), sphingosine-1-phosphate (d18:1), C16 ceramide (d18:1/16:0), C16 dihydroceramide (d18:0/16:0), C18 Ceramide (d18:1/18:0), C16 glucosyl(ß) ceramide (d18:1/16:0), C22 ceramide (d18:1/22:0), C24 ceramide (d18:1/24:0) and C24 dihydroceramide (d18:0/24:0).

### 4.4 Live-cell imaging

Live neuronal imaging experiments were performed, as previously described ^74,75^, by employing a Nikon Eclipse TE300 inverted microscope equipped with a Xenon lamp, a 40× Nikon epifluorescence oil immersion objective (N.A.: 1.3) and a 12-bit Orca CCD camera (Hamamatsu). Alternatively, experiments were performed with a Zeiss Axio Examiner.D1 upright microscope equipped with an Optoscan monochromator (Cairn), a 20x or 40x Zeiss epifluorescence water immersion objective (N.A:1.0), and a 16-bit Evolve 512 EMCCD camera (Photometrics). Images were acquired and stored for offline analysis with Metafluor 7.7 software (Molecular Devices).

### 4.5 [Ca^2+^]_i_ measurements

Measurements of [Ca^2+^]_i_ were performed by employing the ratiometric dye fura-2 or the single wavelength dye fluo-4, as previously described ^74,75^. Briefly, cortical cultures were loaded with fluo-4 AM (3 μM) or fura-2 AM (3 μM) plus 0.1% Pluronic F-127 (ThermoFisher) in a HEPES-buffered saline solution (HCSS) whose composition was: 120 mM NaCl, 5.4 mM KCl, 0.8 mM MgCl_2_, 20 mM HEPES, 15 mM glucose, 1.8 mM CaCl_2_, 10 mM NaOH, and pH 7.4. After 30 min of incubation with the selected dye, cells were washed and incubated in the dark for a further 30 min in HCSS.

Resting [Ca^2+^]_i_ levels were recorded as fura-2 ratios and expressed as % changes compared to vehicle-treated adult cultured neurons. No changes in the time of exposure, gain, electron multiplier settings, filter set, or lamp power occurred during resting [Ca^2+^]_i_ measurements. Pharmacological manipulations were performed by applying drugs at the indicated time points and washed out by employing an automated perfusion system (Biologic). During all the fura-2 [Ca^2+^]_i_ measurements HCSS was supplemented with 200-500 nM TPEN (Merck) to avoid interferences due to other metal ions (i.e. zinc) with [Ca^2 +^]_i_-dependent fluorescence signals ^76,77^.

### 4.6 Spontaneous [Ca^2+^]_i_ spikes analysis

Fluo-4 AM (Ex*λ*:473±20 nm, Em *λ*: 525 ± 25 nm) was employed to measure spontaneous [Ca^2 +^]_i_ transients. Images were acquired at full-frame resolution (512 × 512 pixels; binning 1×) at 1 Hz sampling rate for up to 5 min. Conversely, to evaluate dendritic Ca^2+^ transients, the camera frame was cropped (with a 1x binning) around proximal primary dendritic branches and images acquired at a 10 Hz sampling rate for up to 60 s.

Fluorescence changes of each cell/dendrite were expressed as a percentage of baseline fluorescence (% of basal fluorescence). Spontaneous [Ca^2+^]_i_ transients from vehicle- and L-CS-treated cultures were analyzed with a custom made MATLAB code, as previously described ^28^. For statistical analysis, we took into consideration only fluorescence values 50 % larger than baseline for somata and 25 % larger than baseline for dendrites. Cells that failed to display Ca^2+^ transients when the threshold was set at 25% of the baseline were excluded from the analysis.

### 4.7 Measurements of mitochondrial Δp

Measurements of the mitochondrial Δp were performed as previously described ^74^, with minor modifications. Briefly, cultured cortical neurons were loaded with 50 nM of TMRE for 30 min in culture medium at 37° C. TMRE fluorescence (Ex *λ*: 530 ± 15 nm, Em *λ*: 575–610 nm) changes of each cell (F_x_) were normalized to basal fluorescence intensity (F_0_) obtained by exposing the cells to CCCP (10 µM). Experiments were halted when, after CCCP application, the fluorescence signal was stable for up to 30 s.

### 4.8 Mitochondria morphology analysis

Neurons plated on glass coverslips were washed and loaded for 30 min at room temperature with MitoTracker Green FM (100 nM in HCSS), a mitochondrially-targeted and Δp-insensitive dye. Cells were imaged on the stage of an inverted Zeiss LSM800 Super-resolution confocal microscope equipped with a 488 nm LED-based laser line, a 63x oil immersion objective (N.A.: 1.4), and an Airyscan imaging module. Appropriate emission filter-sets were selected with the built-in Smart Setup function. Laser power was maintained at the minimum (0.65%) to achieve an optimum signal-to-noise ratio and avoid photobleaching. Acquisition parameters (i.e., detector and digital gain) were constant among experimental sessions. Single-plane images were acquired with the Zeiss ZEN proprietary software and stored for offline analysis.

Confocal micrographs were Airyscanned to obtain super-resolution images that were further analyzed with the ImageJ-based MiNA toolkit, following the developer workflow ^26^. Briefly, images were preprocessed (applied algorithms: Unsharp Mask, CLAHE, Median Filtered), then binary transformed and skeletonized to allow parameters generation and recording.

### 4.9 ROS measurements

Evaluation of basal ROS production was performed by employing hydroethidine (HEt; Ex *λ*: 530 ± 15 nm, Em *λ*: 575–610 nm), a fluorescent probe sensitive to ROS. Basal ROS production levels were measured after loading vehicle- or L-CS-treated cultures with HEt (5 µM). Cells were loaded for 2 h at 37 °C to allow dye oxidation by endogenously released ROS. Resting fluorescence signals were recorded and normalized to that of vehicle-treated control cultured neurons.

### 4.10 Western blot analysis

Western blot analysis was performed as previously described ^78,79^. Briefly, control and treated cells were lysed in ice-cold RIPA buffer (phosphate buffer saline pH 7.4 containing 0.5% sodium deoxycholate, 1% Igepal, 0.1% SDS, 5mM EDTA, 1% protease and phosphatase inhibitor cocktails). Protein lysates (20 μg) were separated on 8–14% SDS–polyacrylamide gel and electroblotted onto a polyvinyldifluoride membrane (PVDF). Nonspecific binding sites were blocked by 5% non-fat dry milk (Bio-Rad Laboratories) in Tris buffered saline (TBS: 20mMTris– HCl, pH 7,4, containing 150mM NaCl) for 30 min at RT. Membranes were then incubated overnight at 4°C with the following primary antibodies, diluted with TBS containing 0,1% Tween 20 (TBS-T) and 5% non-fat dry milk: rabbit HRP-conjugated actin 1:10000 (Cell Signaling); rabbit p-JNK and JNK 1:200 (Santa Cruz); p-ERK1,2 1:200 (Santa Cruz); rabbit p-Akt 1:1000 (Cell Signaling); Erk1,2 1:200 (Santa Cruz); rabbit p-P38 and rabbit P38 1:1000 (Cell Signaling); rabbit P53 1:1000 (Cell Signaling); rabbit p-ERK5 and rabbit ERK5 1:1000 (Cell Signaling). As secondary antibodies, peroxidase conjugated anti-rabbit or anti mouse IgG (1:10000; Vector Laboratories) were used. Immunoreactive bands were visualized by ECL (Bio-Rad Laboratories), according to the manufacturer’s instructions. The relative densities of the immunoreactive bands were determined and normalized with respect to Actin, using ImageJ software. Values were given as relative units (RU).

### 4.11 Data and statistical analysis

Data are represented as box plots. Center lines and boxes represent medians and means, respectively. Box limits indicate 25^th^ and 75^th^ percentiles, and whiskers extend 1.5 times the interquartile range from the 25^th^ and 75^th^ percentiles ^80^. No statistical methods were used to predetermine the sample size. Statistical analysis was performed by two-way ANOVA (cell culture age vs. treatment) followed by Tukey’s post-hoc test. Differences in ceramide levels and dendritic Ca^2+^ transients were analyzed by unpaired Student’s t-test. Experimenters were not blinded to treatment allocation. By conventional criteria, P values are represented as * for P ≤ 0.05 and ** for P ≤ 0.01.

## Acknowledgments

The authors thank all the members of the Molecular Neurology Unit for helpful discussions. SLS is supported by research grants from the Italian Department of Health (RF-2013–02358785 and NET-2011-02346784-1), from the AIRAlzh Onlus (ANCC-COOP), from the Alzheimer’s Association - Part the Cloud: Translational Research Funding for Alzheimer’s Disease (18PTC-19-602325) and the Alzheimer’s Association - GAAIN Exploration to Evaluate Novel Alzheimer’s Queries (GEENA-Q-19-596282). AG is supported by the European Union’s Horizon 2020 research and innovation programme under the Marie Skłodowska-Curie grant agreement iMIND – No. 841665.

## Author contribution

SLS, DP, and AG conceived and designed the study. AG performed and analyzed all the imaging experiments. MB helped with neuronal culturing. RN and NM analyzed spontaneous Ca^2+^ imaging experiments. IC and PdB performed and analyzed ceramide levels. VC and AC performed and analyzed WB data. AG and SLS analyzed and interpreted the data. MO and DP reviewed and provided a critique of the study design and writing of the manuscript. AG and SLS wrote the manuscript.

---

**Supplementary Figure 1.**
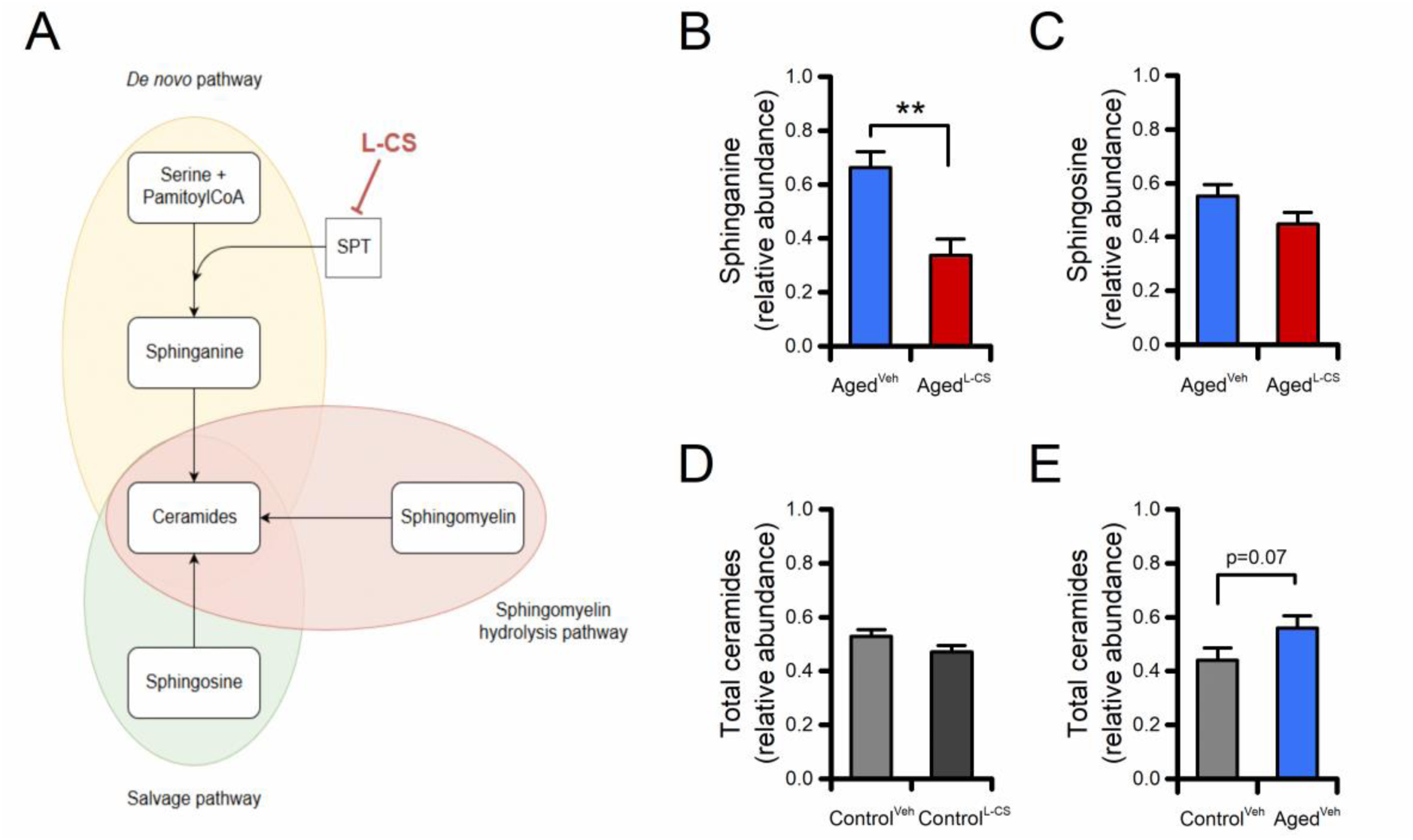
Effects of aging and L-CS on *de novo* ceramide biosynthesis. (A) The pictogram illustrates a simplified version of the steps involved in the three ceramide biosynthetic pathways. Please, note that L-CS acts by inhibiting SPT in the *de novo* pathway. (B) Bar graphs depict the relative abundance of sphinganine in vehicle- and L-CS-treated aged neurons (n=3). (C) Bar graphs depict the relative abundance of sphingosine in vehicle- and L-CS-treated aged neurons (n=3). (D) Bar graphs depict the relative abundance of ceramides in vehicle- and L-CS-treated control neurons (n=3). (E) Bar graphs depict the relative abundance of ceramides in vehicle-treated control and aged neurons (n=3). Means were compared by unpaired Student t-test. * * indicates p<0.01.

**Supplementary Figure 2.**
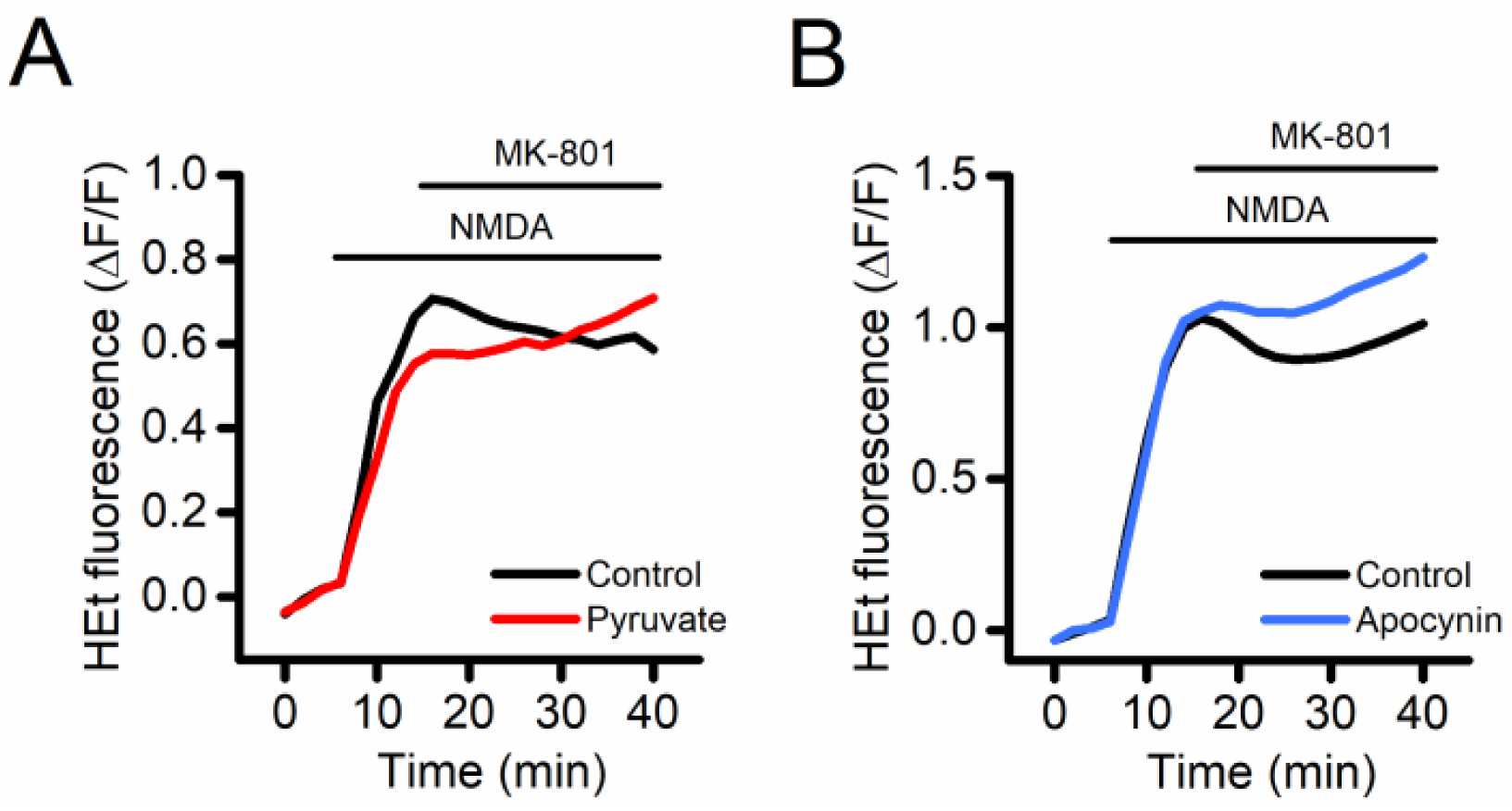
Mitochondria are the primary source of ROS in our cortical cultures. To evaluate the primary source of ROS in our system we performed pharmacological manipulations aimed at selectively observing ROS of mitochondrial and non-mitochondrial origin. HEt-loaded control cortical neurons were challenged with NMDA + glycine (50 µM + 10 µM), a maneuver that triggers a robust generation of ROS from both mitochondrial and non-mitochondrial sources ^81,82^. After 5 minutes, NMDA receptor (NMDAR) overactivation was halted by bath application of the non-competitive NMDAR antagonist MK-801 (10 µM). In (A), traces depict NMDA-driven ROS generation in control neurons (black trace) and in cells bathed and challenged in a solution in which glucose was replaced with pyruvate (15 mM), an established paradigm aimed at promoting ROS generation only from mitochondria. No differences were observed between the two conditions. In (B), neurons were challenged with (blue trace) or without (black trace) apocynin (500 µM), an inhibitor of NADPH oxidase activity. No differences were observed between the two conditions. Collectively, these findings indicate that mitochondria are the primary source of ROS in our cortical cultures.

**Supplementary table 1.** Data related to mitochondrial morphology analysis as shown in Fig. 3.

|  | **Control<sup>Veh</sup>** | **Control<sup>L-CS</sup>** | **Aged<sup>Veh</sup>** | **Aged<sup>L-CS</sup>** | **P** |
| --- | --- | --- | --- | --- | --- |
| **Individuals (*n*)** | 191.00 ± 46.27 | 168.4 ± 34.11 | 251.00 ± 33.49 | 217.80 ± 54.14 | >0.05 |
| **Networks (*n*)** | 64.25 ± 16.28 | 38.6 ± 11.69 | 78.16 ± 9.81 | 71.2 ± 15.61 | >0.05 |
| **Mean Branch Length (μm)** | 0.602 ± 0.020 | 0.579 ± 0.012 | 0.560 ± 0.016 | 0.547 ± 0.010 | >0.05 |
| **Median Branch Length (μm)** | 0.456 ± 0.014 | 0.467 ± 0.013 | 0.435 ± 0.014 | 0.417 ± 0.007 | >0.05 |
| **Length Standard Deviation (μm)** | 0.531 ± 0.035 | 0.464 ± 0.015 | 0.486 ± 0.017 | 0.490 ± 0.008 | >0.05 |
| **Median Network Size (*n* of branches)** | 4.25 ± 0.47 | 4.40 ± 0.40 | 4.41 ± 0.55 | 4.10 ± 0.45 | >0.05 |
| **Mitochondrial Footprint (μm<sup>2</sup>)** | 139.19 ± 30.06 | 241.26 ± 47.43 | 189.39 ± 27.35 | 153.51 ± 24.35 | >0.05 |

